# The Feeding Connectome: Convergence of Monosynaptic and Polysynaptic Sensory Paths onto Common Motor Outputs

**DOI:** 10.1101/368878

**Authors:** Anton Miroschnikow, Philipp Schlegel, Andreas Schoofs, Sebastian Hückesfeld, Feng Li, Casey M Schneider-Mizell, Richard D Fetter, James Truman, Albert Cardona, Michael J Pankratz

## Abstract

Little is known about the organization of central circuits by which external and internal sensory inputs act on motor outputs to regulate fundamental behaviors such as feeding. We reconstructed, from a whole CNS EM volume, the synaptic map of input and output neurons that underlie food intake behavior of *Drosophila* larvae. The input neurons originate from enteric, pharyngeal and external sensory organs and converge onto seven distinct sensory synaptic compartments within the CNS, as defined by distribution patterns of their presynaptic sites. The output neurons consist of pharyngeal motor neurons, serotonergic modulatory neurons, and neuroendocrine neurons that target the ring gland, a key endocrine organ. Monosynaptic connections from a set of sensory synaptic compartments cover the motor and endocrine targets in overlapping domains. Polysynaptic routes can be superimposed on top of the monosynaptic connections, resulting in divergent sensory paths that converge on common motor outputs. A completely different set of sensory compartments is connected to the mushroom body calyx of the memory circuits. Our results illustrate a circuit architecture in which monosynaptic and multisynaptic connections from sensory inputs traverse onto output neurons via a series of converging paths.

## Introduction

Motor outputs of a nervous system can be broadly defined into those carried out by the muscles to produce movements and by the neuroendocrine glands for secretion (*Shepherd 1987*). Both of these behavioral and physiological events are regulated by a network of motor neurons, interneurons and sensory neurons, and a major open question is how one neural path is selected from multiple possible paths to produce a desired motor output (*Grillner et al. 2005*). Nervous system complexity and tool availability have strongly dictated the type of experimental system and analysis that can be used to address this issue, such as a focus on a particular organism, behavior or type of neuron. In this context, the detailed illustrations of different parts of nervous systems at neuronal level as pioneered by Cajal, to the first complete description of a nervous system wiring diagram at synaptic level for *C. elegans*, demonstrate the power of systematic neuroanatomical analysis in providing a foundation and guide for studying nervous system function (*Ramon y Cajal 1894; White et al. 1986*). However, the technical challenges posed by such analysis have limited the type of organisms for which synaptic resolution mapping can be performed at the scale of an entire nervous system (*Swanson and Lichtman 2016; Schlegel et al. 2017; Kornfeld and Denk 2018*).

Analysis of the neural circuits that mediate food intake in the *Drosophila* larvae offers numerous advantages in meeting the challenge of neuroanatomical mapping at a whole brain level, and combining it with the ability to perform behavioral and physiological experiments. The muscle system that generates the different movements necessary for transporting food from the pharynx to the esophagus, as well as the endocrine system responsible for secreting various hormones for metabolism and growth, have both been well described (*Kühn 1971; Siegmund and Korge 2001; Buch and Pankratz 2009; Schoofs et al. 2010*). These are also complemented by the analysis of feeding behavior in adult flies (*Gelperin 1971; Dethier 1976; McKellar 2016*). Although there is broad knowledge at the morphological level on the organs underlying larval feeding behavior and physiology, as well as on the nerves innervating them in the periphery (*Schoofs et al. 2010, 2014b*), the central connectivity of the afferent and efferent neurons within these nerves are largely unknown. At the same time, advances in the EM reconstruction of an entire CNS of a first instar larva (*Ohyama et al. 2015; Schlegel et al. 2016; Schneider-Mizell et al. 2016; Berck et al. 2016; Eichler et al. 2017; Gerhard et al. 2017*) (summarized in *Kornfeld & Denk, 2018*) offers an opportunity to elucidate an animals’ feeding system on a brain-wide scale and at synaptic resolution. As part of this community effort, we recently performed an integrated analysis of fast synaptic and neuropeptide receptor connections for an identified cluster of 20 interneurons that express the neuropeptide hugin, a homolog of the mammalian neuropeptide neuromedin U, and which regulates food intake behavior (*Melcher et al. 2006; Schoofs et al. 2014a; Schlegel et al. 2016*). This analysis showed that the class of hugin neurons modulating food intake receives direct synaptic inputs from a specific group of sensory neurons, and in turn, makes mono-synaptic contacts to output neuroendocrine cells. The study not only provided a starting point for a combined approach to studying synaptic and neuropeptidergic circuits (*Diao et al. 2017; Williams et al. 2017*), but a basis for a comprehensive mapping of the sensory and motor neurons that innervate the major feeding and endocrine organs.

Feeding is one of the most universal and important activities that animals engage in. Despite large differences in the morphology of the external feeding organs, the internal gut structures are quite similar across different animals (*Campbell 1990*); indeed, even within closely related species, there can be large differences in the external organs that detect and gather food, whereas the internal organs that transport food through the alimentary canal are much more similar. Recent studies have also pointed out the functional similarities between the subesophageal zone in insects and the brainstem in vertebrates for regulating feeding behavior (*Schoofs et al. 2014a; Yapici et al. 2016; McKellar 2016*). In mammals, the different cranial nerves from the medulla innervate distinct muscles and glands of the foregut (***Figure 1A***). For example, the VIIth cranial nerve (facial nerve) carries taste sensory information from anterior 2/3 of the tongue, and innervates the salivary glands, and lip and facial muscles. The IXth cranial nerve (glossopharyngeal nerve) receives taste inputs from the posterior 1/3 of the tongue, and innervates the salivary glands and pharynx muscles. The Xth cranial nerve (vagus nerve) receives majority of the sensory inputs from the enteric nervous system of the gut, and innervates pharynx and esophagus muscles. The XIth cranial nerve (spinal accessory nerve) and the XIIth cranial nerve (hypoglossal nerve) are thought to carry strictly motor information which innervate the pharynx and neck muscles, and the tongue muscles (*Cordes 2001; Simon et al. 2006*). The distinct cranial nerves project onto topographically distinct areas in the medulla of the brainstem (***Figure 1A***). We also note that olfactory information is carried by cranial nerve I, a strictly sensory nerve that projects to the olfactory bulb (OB), an area topographically distinct from the brainstem. In addition, there are direct neuronal connections between the brainstem and the hypothalamus, the key neuroendocrine center of vertebrates (*D’Agostino et al. 2016; Liu et al. 2017*).

**Figure 1.**
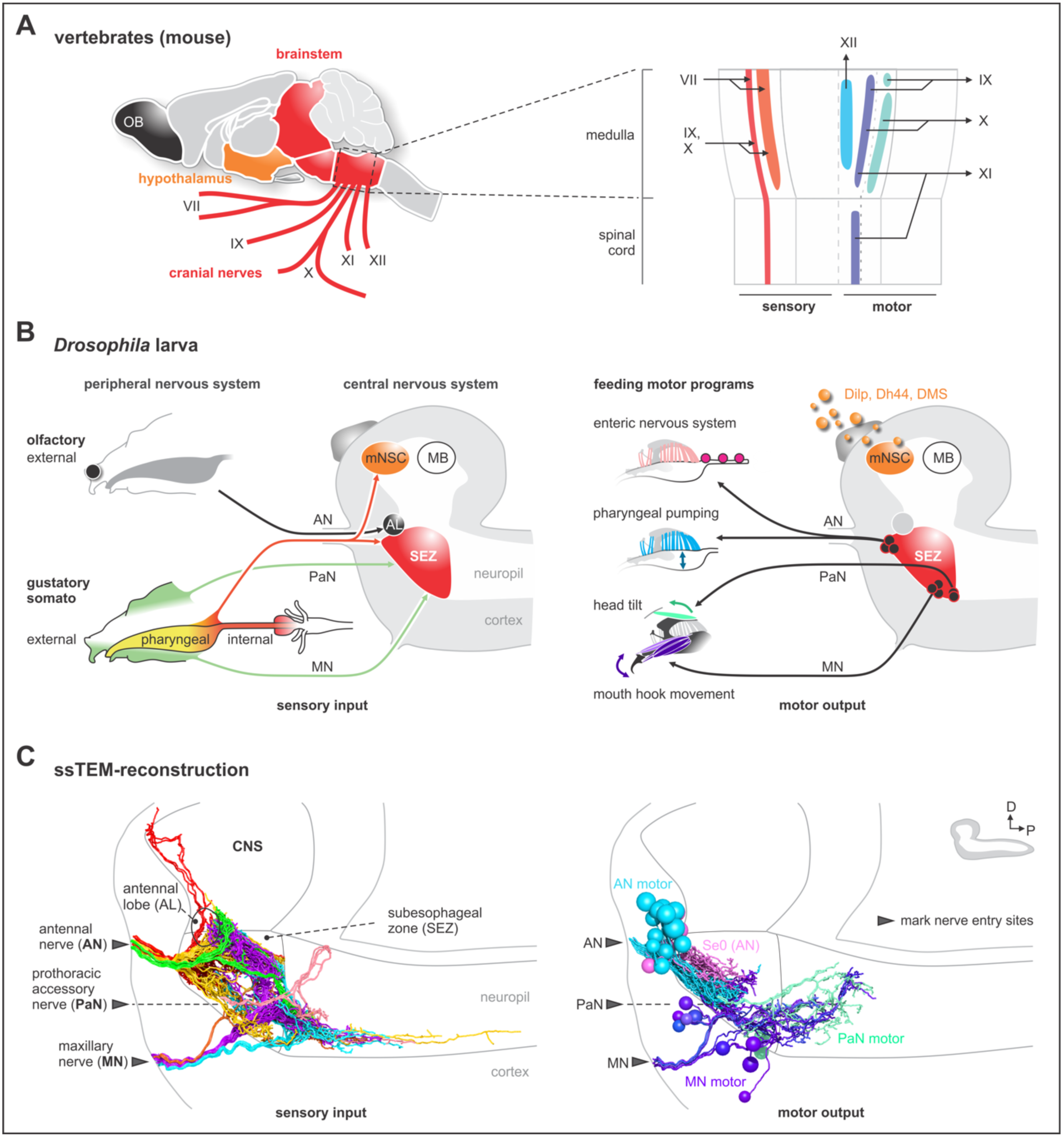
EM reconstruction of the pharyngeal nerves of *Drosophila* larva. **(A)** Left: schematic diagram shows a lateral view of an adult mouse brain and the broad organization of different cranial nerves targeting the medulla of the brainstem. Right: topological chart of the medulla and part of the spinal cord. Primary sensory and primary motor nuclei are shown on the left and on the right, respectively. **(B)** Schematic overview of external, pharyngeal and internal sensory systems targeting the subesophageal zone (SEZ), median neurosecretory cells (mNSCs) and the antennal lobe (AL) in *Drosophila* (left panel). Schematic overview of central output neurons targeting feeding related muscles (right panel). Median neurosecretory cells (mNSCs) target neuroendocrine organ and the periphery, by releasing neuropeptides such as Dilp, Dh44 and DMS. The mushroom body (MB), a learning and memory center, serves as a landmark. **(C)** EM reconstruction of pharyngeal sensory input (left panel). Sensory neurons enter the brain via the antennal nerve (AN), maxillary nerve (MN) and prothoracic accessory nerve (PaN), and cover large parts of the SEZ (left panel). Arrowheads mark respective nerve entry site. Two of the AN sensory projections (per side) extend into the protocerebrum. EM reconstruction of pharyngeal motor output (right panel). AN motor neurons and serotonergic output neurons (Se0) leave the CNS via the antennal nerve (AN) and innervate the cibarial dilator musculature (for pharyngeal pumping) and part of the esophagus and the enteric nervous system. MN motor neurons leave the CNS via the maxillary nerve (MN) and innervate mouth hook elevator and depressor, labial retractor and salivary gland ductus opener. PaN motor neurons leave the CNS via the prothoracic accessory nerve (PaN) and innervate the dorsal protractor (for head tilt movements). All neurons are colored based on their morphological class. See ***Figure 1–figure supplement 1-4*** for detailed anatomy and description of morphological clustering.

Analogously, distinct pharyngeal nerves of the *Drosophila* larva are connected to the subesophageal zone (SEZ), and also carry sensory and motor information that regulate different parts of the body (***Figure 1B***). The AN (antennal nerve) carries sensory information from the olfactory, pharyngeal and internal organs, and innervates the pharyngeal muscles for pumping in food.

The neurons that innervate the major endocrine center and the enteric nervous system also project through the AN. Note also that the olfactory sensory organs project to the antennal lobe (AL), which abuts the SEZ yet is topographically separate. The MN (maxillary nerve) carries external and pharyngeal sensory information, and innervates the mouth hooks, whose movements are involved in both feeding and locomotion. The PaN (prothoracic accessory nerve) carries external sensory information from the upper head region, and innervates the muscles involved in head tilting (see ***Figure 1–figure supplement 1*** for anatomical details). Furthermore, the SEZ has direct connections to median neurosecretory cells and the ring gland. In sum, although a large body of knowledge exists on the gross anatomy of the nerves that target the feeding organs in vertebrates and invertebrates, the synaptic pathways within the brain that interconnect the sensory inputs and motor outputs of the individual nerves remain to be elucidated.

In this paper, we have reconstructed all sensory and motor neurons of the three pharyngeal nerves that underlie the feeding motor program of *Drosophila* larvae. The activity of these nerves has previously been shown to be sufficient for generating the feeding motor pattern in isolated nervous system preparations, and that the central pattern generators (CPGs) for food intake lie in the SEZ (*Schoofs et al. 2010; Hückesfeld et al. 2015*). We then identified all monosynaptic connections between the sensory and motor neurons, thus providing a full monosynaptic reflex circuit for food intake. We also mapped polysynaptic pathways that are integrated onto the monosynaptic reflex circuits. In addition, we mapped the multisynaptic non-olfactory neuron connections from the sensory neurons to the mushroom body memory circuit (*Eichler et al. 2017*), and show that these are different from those involved in monosynaptic reflex circuits. Reflex circuits can be seen to represent the simplest synaptic architecture in the nervous system, as formulated by Charles Sherrington (*Sherrington 1906*). Anatomical reconstructions of monosynaptic and poly-synaptic reflex circuits can also be seen in the works of Cajal (*Ramon y Cajal 1894; Swanson 2000*). We propose a model of how different mono- and polysynaptic pathways can be traversed from a set of sensory neurons to specific output neurons, which has relevance for understanding the mechanisms of action selection.

## Results

### EM reconstruction of the pharyngeal nerves

We reconstructed all axons within the three pharyngeal nerves into the CNS using a ssTEM volume of an entire larval CNS (*Ohyama et al. 2015*) (***Figure 1C***). The sensory projections were those that ended blindly, whereas the motor neurons were those with somata in the CNS. For sensory inputs, a regionalization of the target areas can already be seen, reflecting the fact that the nerves are fusions of different axon bundles that arise during embryonic development (*Hartenstein et al. 2017; Kendroud et al. 2017*). For example, only the AN has sensory projections that extend into the protocerebrum, whereas a major part of the MN sensory projections extend into the ventral nerve cord. For motor neurons, the somas from the different nerves also occupy distinct regions within the SEZ (***Figure 1C***; ***Figure 1–figure supplement 1-4*** for details on individual nerves and bundles).

### Topographical patterns of sensory and motor synaptic compartments in the CNS

We next annotated all pre- and postsynaptic sites of all sensory projections and clustered them based on synapse similarity (***Figure 2A***). This revealed 7 topographically distinct compartments in the CNS (***Figure 2B***; ***Figure 2–figure supplements 1-2***). These compartments differ in the number of sensory neurons as well as in the identity of the pharyngeal nerve that gives rise to them (***Figure 2–figure supplements 3-5***). For example, the ACa (“anterior part of the Anterior Central sensory compartment”) comprises 30 neurons that are exclusively derived from the AN; by contrast, the VM (“Ventromedial sensory compartment”) comprises 102 neurons that derive from all three pharyngeal nerves. For endocrine output neurons, we previously reconstructed the three neurosecretory cell clusters of the pars intercerebralis that innervate the major endocrine organ of *Drosophila* larvae (the ring gland). These express the neuropeptides Dilps (*Drosophila* insulin-like peptides), DH44 (diuretic hormone, a corticotropin regulating hormone homolog) and DMS (*Dro*myosuppressin), and receive monosynaptic inputs from the sensory system (***Figure 2C,D***) (*Schlegel et al. 2016*). We now identify here all pre- and post-synaptic sites of all the motor neurons of the different pharyngeal nerves (***Figure 2C***). This includes a special class of four serotonergic neurons (the Se0 cluster) that project to the entire enteric nervous system (*Huser et al. 2012; Schoofs et al. 2014b; Shimada-Niwa and Niwa 2014*). These four serotonergic neurons can be further divided into one that projects anteriorly to the pharynx (Se0ph), and three that project posteriorly towards the enteric nervous system (Se0ens) (***Figure 2–figure supplement 6***).

**Figure 2.**
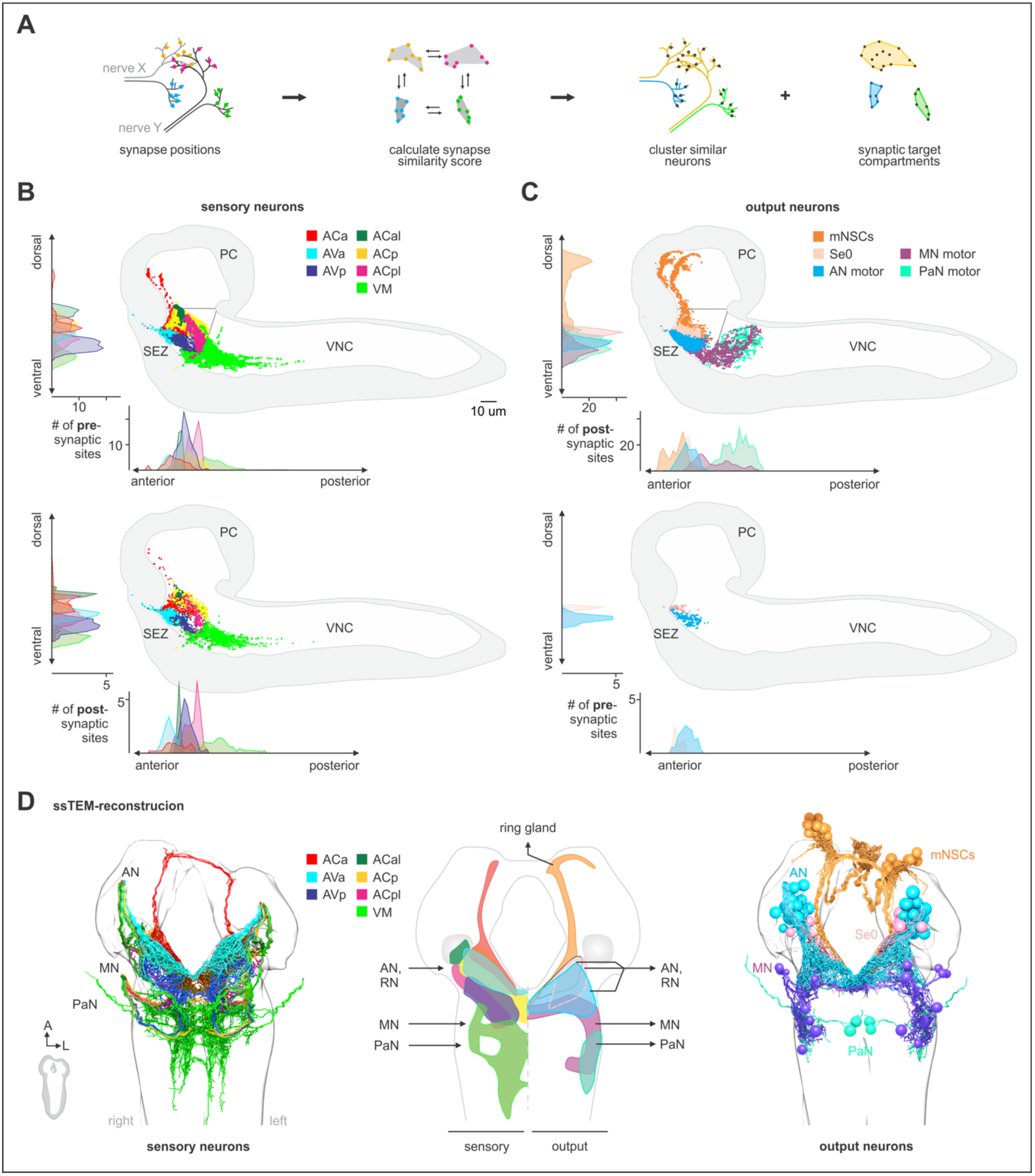
Spatially segregated central axonal projections of sensory neurons. **(A)** Calculation of pairwise synapse similarity score for all non-olfactory sensory neurons. **(B)** Spatial distribution of synaptic sites for all sensory neuron cluster. Hierarchical clustering based on synapse similarity score revealed 7 distinct (non-overlapping) areas of sensory convergence within the SEZ: the anterior part of the Anterior Central sensory compartment (ACa), anterior part of the Anterior Ventral sensory compartment (AVa), posterior part of the Anterior Ventral sensory compartment (AVp), posterior part of the Anterior Central sensory compartment (ACp), anterior-lateral part of the Anterior Central sensory compartment (ACal), posterior-lateral part of the Anterior Central sensory compartment (ACpl) and Ventromedial sensory compartment (VM). Convergence zones are targeted by varying numbers of sensory neurons but are consistent across hemispheres. Each dot represents a single synaptic site. Graphs show distribution along dorsal-ventral and anterior-posterior axis of the CNS. **(C)** Spatial distribution of synaptic sites for all endocrine and motor neuron classes. Each dot represents a single synaptic site. Graphs show distribution along dorsalventral and anterior-posterior axis of the CNS. **(D)** EM reconstruction of clustered sensory neurons (left). EM reconstruction of all output neuron classes (right). Summarizing representation of glomerular-like sensory compartments and motor compartments within the SEZ (middle panel). See ***Figure 2–figure supplement 1-5*** for detailed description of clustering and sensory region composition.

A schematic summary of the pre- and post-synaptic compartments of the input and output neurons, along with their projection regions, is shown in ***Figure 2D***. Taken together, these data define all sensory input convergence zones and motor output compartments of the three pharyngeal nerves underlying feeding motor program at synaptic resolution.

### Axo-dendritic connections from sensory to output motor neurons

Having annotated all central synapses of in- and output neurons, we surprisingly found the most basic element of circuit architecture: direct monosynaptic connections between input and output neurons (***Figure 3A***). The vast majority of the monosynaptic connections are made from anterior 3 of the 7 sensory compartments (ACa, AVa, AVp; ***Figure 3B,C***; ***Figure3–figure supplement 2***): around 90% of the neurons in these 3 compartments make monosynaptic contacts.

Importantly, in- and outputs compartments do not perfectly overlap. As a consequence, we find single input compartments to make monosynaptic connections to neurons in overlapping output compartments, as one progresses from neuroendocrine (mNSCs), serotonergic neuromodulatory (Se0), and pharyngeal motor neurons (***Figure 3D***; ***Figure3–figure supplement 2-5***): ACa inputs onto the neuroendocrine and Se0 neurons; AVa onto neuroendocrine, Se0 and AN motor neurons; AVp onto Se0 and AN motor neurons; VM onto MN and PaN motor neurons. Thus, the monosynaptic connections essentially cover all output neurons in contiguous, overlapping domains. When viewed from the sensory neuron side, a small percentage (less than 5% of synapses) makes monosynaptic contacts (***Figure 3E*** left panel); however, when viewed from the output neuron side, the percentage of monosynaptic inputs they receive are substantial (***Figure 3E*** right panel). For example, around 40% of all input synapses onto the serotonergic Se0 neurons, and between 10-25% of all mNSC input synapses, are from sensory neurons (***Figure 3F***; ***Figure 3–figure supplement 2-5*)**. In sum, these results indicate that the monosynaptic connections between sensory and motor output neurons form a special class with which a core or an “elemental” feeding circuit can be constructed (***Figure 3C***; ***Figure 3–figure supplement 1***).

**Figure 3.**
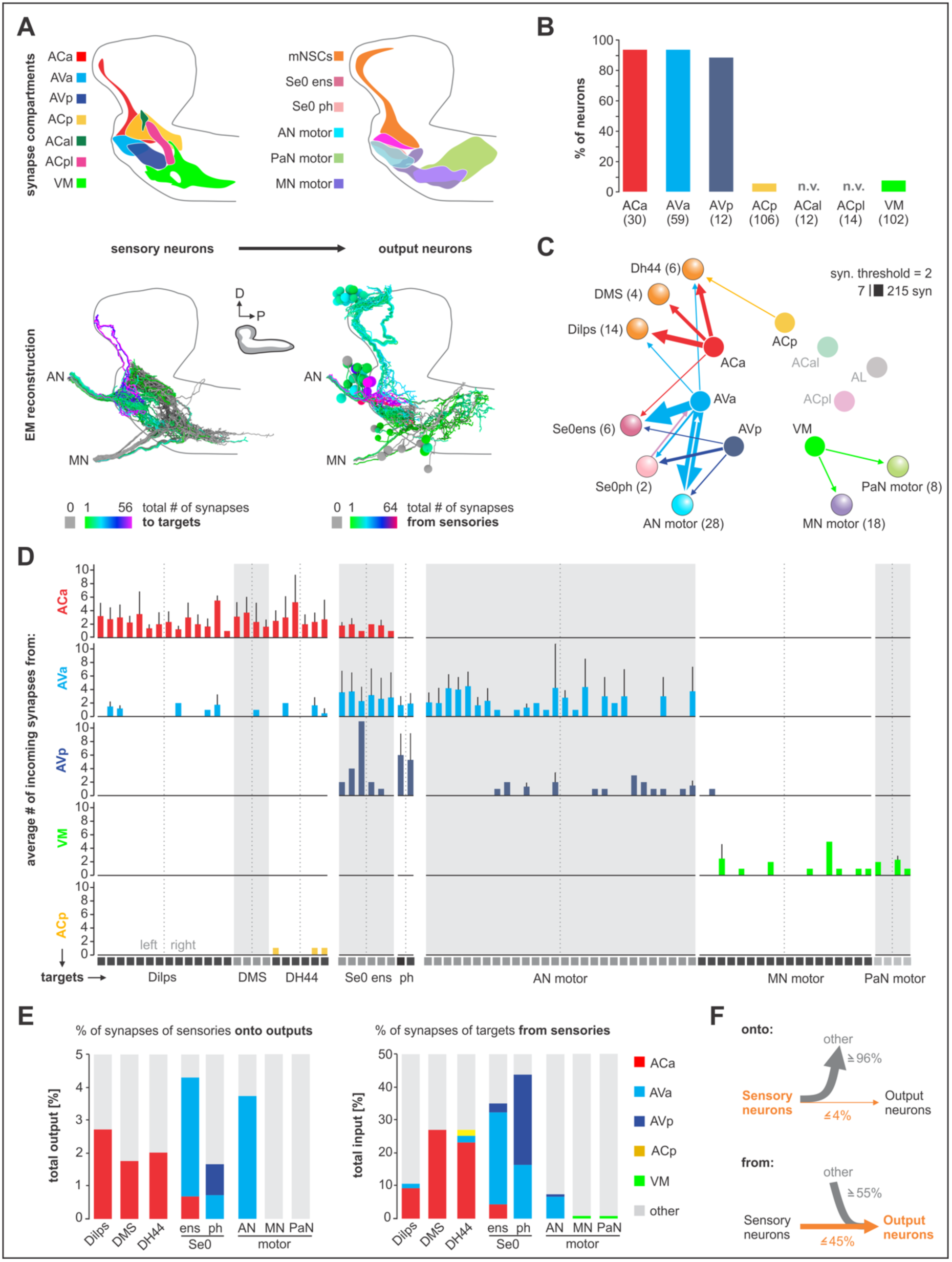
Monosynaptic circuits between sensory and output neurons. **(A)** Lateral schematics of the presynaptic sensory compartments and postsynaptic terminals for each output neuron type (upper panel). EM reconstruction of respective neurons (lower panel). Left: sensory neurons are color-coded based on total number of synapses to their monosynaptic targets. Right: output neurons are color-coded based on total number of synapses from sensory neurons. Lateral views show neurons of the right side. **(B)** Percentage of sensory neurons of the respective sensory compartment forming monosynaptic circuits. About 90% of all sensory neurons of ACa, AVa, AVp are part of monosynaptic circuits; in contrast, ACp, ACal, ACpl and VM show little to none. **(C)** Connectivity diagram of axo-dendritic connections between sensory and output neurons. Circles represent previously defined sensory and motor compartments (***Figure 2***). Sensory compartments with no monosynaptic reflex connections are faded. **(D)** Connectivity between presynaptic sensory neurons (ACa, AVa, AVp, ACp and VM) and postsynaptic output targets. Each column across all five sensory compartments represents a postsynaptic partner of the sensory neurons. Whiskers represent standard deviation. **(E)** Left: percentage of synapses of sensory neurons to monosynaptic targets. Right: percentage of synapses of output neurons from sensory neurons. **(F)** Summarizing representation of monosynaptic input-to-output ratio viewed from the sensory neuron side (top) or from the output neuron side (bottom). See ***Figure 3— figure supplement 1-5*** for detailed connectivity.

### Axo-axonic connections between sensory neurons

Unexpectedly we found a high number of synaptic connections between the sensory projections within the CNS. This is in contrast to the well characterized olfactory sensory neurons that project onto the antennal lobe (AL) (***Figure 4A***). For example, at a threshold of two synapses the AL has none, whereas 50% of the ACa neurons have above threshold inter-sensory connections. The majority of the inter-sensory connections were between neurons of the same synaptic target compartment (***Figure 4B***), which underscores the clear-cut boundaries between the sensory compartments; these connections are made both in an hierarchical manner as well as reciprocally, suggesting that sensory information processing is occurring already at an inter-sensory level in the brain (***Figure 4–figure supplement 1***). Viewed from output synapses of the sensory neurons, the percentage of sensory synapses connecting to other sensory neurons are small relative to total sensory outputs (less than 2% of 73,000 synapses); however viewed from input side of the sensory neurons, a high percentage (e.g., 45% for ACa) of their total synaptic inputs originate from other sensory neurons (***Figure 4C,D***). We also note that sensory neurons from ACp and VM have inter-sensory connections even between neurons of different nerves (***Figure 4–figure supplement 1-2***), indicating integration of sensory information from different body regions at the sensory neuron level.

**Figure 4.**
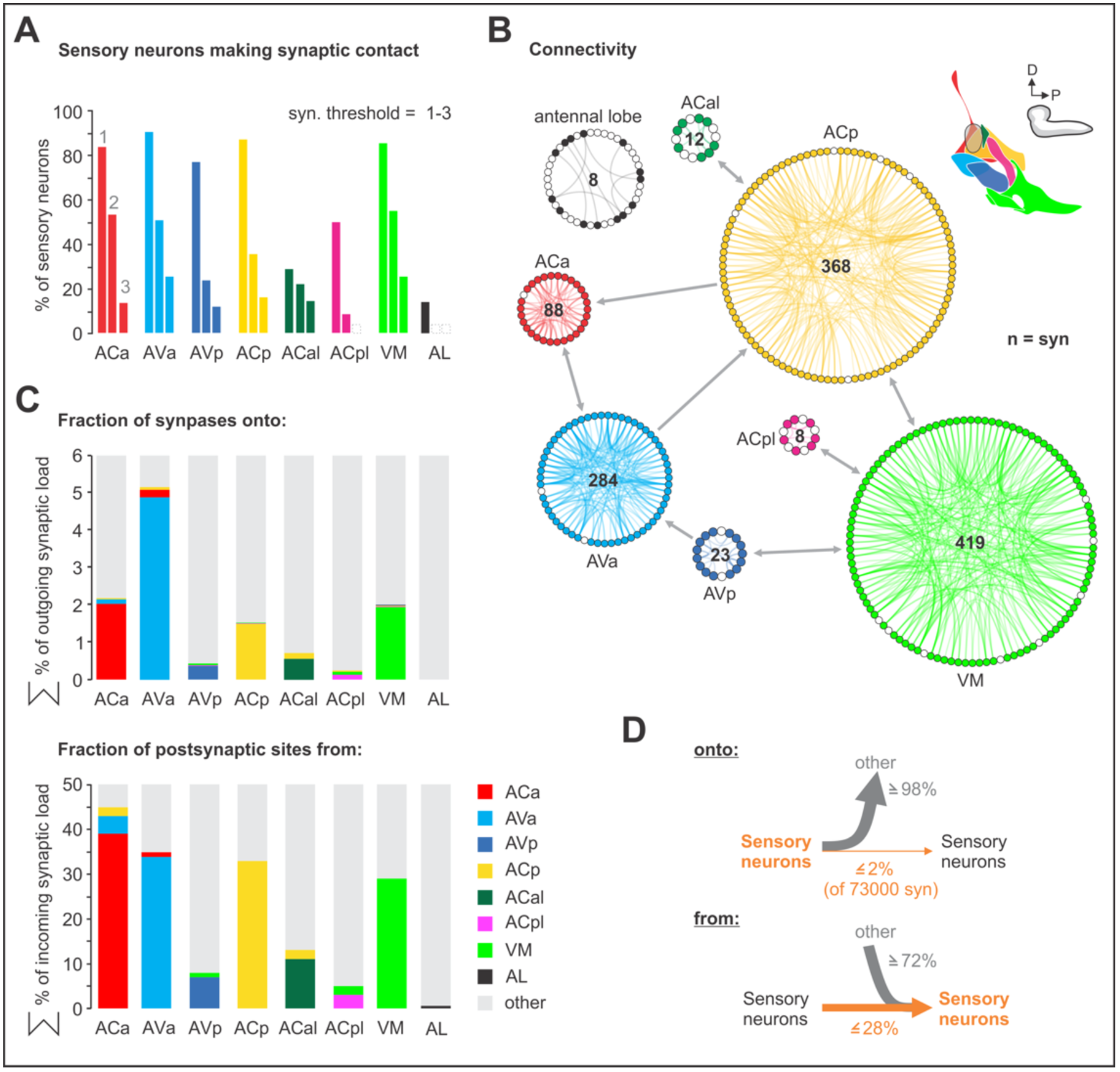
Sensory-sensory communication. **(A)** Percentage of sensory neurons of sensory clusters involved in intra cluster sensory connections. Around 80% of all sensory neurons in ACa, AVa, AVp, ACp and VM form intra sensory connections. ACal and ACpl have the lowest number of neurons and also show low number of intra sensory contacts. **(B)** Connectivity diagram of axo-axonic connections between sensory neurons. The circular wheel arrangements represent previously defined sensory compartments (see ***Figure 2***). Each small circle within a wheel represents a single neuron. Gray links show inter-cluster connections (max. 10 synapses in one direction). Note that sensory to sensory contacts are made mainly between sensory neurons of the same class, not between classes. **(C)** Percentage of synapses of sensory neurons from and onto sensory neurons. **(D)** Summarizing representation of axo-axonic contact input-to-output ratio viewed from the presynaptic neuron side (top) or from the postsynaptic neuron side (bottom). See ***Figure 4–figure supplement 1-2*** for detailed connectivity.

### Mapping peripheral origins of monosynaptic circuits

We next investigated the peripheral origins of the sensory neurons that comprise the different synaptic compartments. This was accomplished by using various sensory receptor Gal4 lines to follow the projections from the sensory organs into the CNS. The mapping was aided by the fact that the pharyngeal projections enter the SEZ in distinct bundles that can be observed in both light and EM microscopic sections (***Figure 5A,B***). The AN and the MN each have three bundles (these nerves are formed by fusion of several axon bundles during development (*Hartenstein et al. 2017*), whereas the PaN has just one. The well characterized projections from the external olfactory organ (DOG) to the antennal lobe (AL), for example, use one of the bundles in the AN (Bundle 3 of the AN). ***Figure 5B*** illustrates the basic strategy, using two of the gustatory receptors (GRs) to follow the projections from the enteric nervous system into the CNS. This analysis, denoting the receptor line used and their expression in the sensory organs and the axon bundles of each pharyngeal nerve, is summarized in ***Figure 5C*** (***Figure 5–figure supplement 1-9*** for detailed stainings). These results were then used to determine the peripheral origin (enteric/internal, pharyngeal, external) of the sensory neurons that comprise the 7 synaptic compartments defined earlier (***Figure 5C,D***). This revealed a wide spectrum in compartment composition. For example, the ACa is derived 100% from the enteric nervous system, while the AVa is 93 % enteric; these are the only two sensory compartments with enteric origin. As a comparison, the antennal lobe (AL) is derived 100% from a single external sensory organ, the dorsal organ. Interestingly, the topographical location of the sensory compartments within the CNS broadly mirrors in a concentric manner the peripheral origins from which they derive: the inner-most enteric organs project to the anterior most region, the pharyngeal sensory organs project to the middle region, while the most external organs project to the outer-most region (***Figure 5E***). Recent light microscopy study on the projections of somatosensory neurons onto the adult brain also showed topographically separate target areas in the brain (*Tsubouchi et al. 2017*).

**Figure 5.**
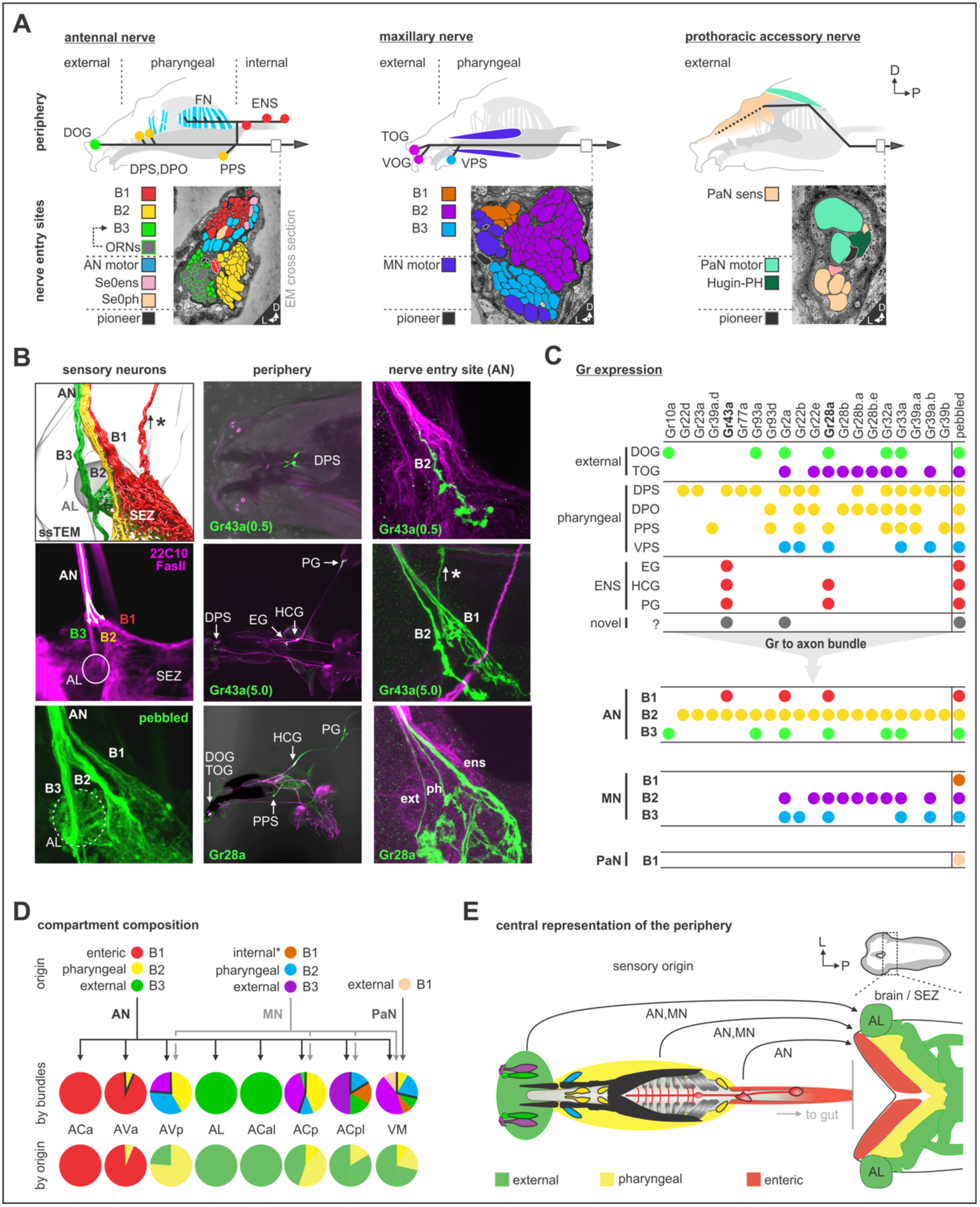
Mapping peripheral origin of sensory neurons. **(A)** Origins and targets of feeding related sensory and motor neurons. The AN comprises motor axons innervating the cibarial dilator muscles (blue striped region) and sensory axons from the dorsal organ ganglion (DOG), pharyngeal sensilla (DPS, DPO, PPS), frontal nerve (FN) and enteric nervous system (ENS). The MN comprises motor axons innervating the mouth hook elevator and depressor (in purple), labial retractor and salivary gland ductus opener; and sensory axons from the terminal organ ganglion (TOG), ventral organ ganglion (VOG) and pharyngeal sensilla (VPS). The PaN comprises motor axons innervating the dorsal protractor (in green), and sensory neurons with an hypothesized origin in the anterior pharyngeal region (in beige). EM cross section of the right AN, MN and PaN at nerve entry site (lower panels). Neuronal profiles of all neurons are colored based on their morphological class and origin. **(B)** Mapping of Gr28a and Gr43a gustatory receptor neuron projection through distinct bundles of the AN from the enteric nervous system. Pebbled-Gal4 was used as a pan-sensory neuronal marker to shows expression in all 3 bundles of the AN. Asterisk marks sensory projections into the protocerebrum. **(C)** Summary table of selected Gr expression patterns from the peripheral origin (sensory organs and ganglia), and their expression in respective nerve entry site. Note that Gr28a and Gr43a show expression in the ENS (EG, esophageal ganglion; HCG, hypocerebral ganglion; PG, proventricular ganglion), which results in projections through bundle 1 (B1). **(D)** Sensory compartment composition by peripheral origin. ACa, ACal and AL each derive from a single sensory origin. In contrast, AVa, AVp, ACp, ACpl and VM integrate several sensory origins. Percentage compartment composition is shown by nerve bundles and by origin (enteric, pharyngeal, external). **(E)** Somatotopic arrangement of sensory axon in the brain and SEZ, showing a layered arrangement that mirrors the antero-posterior layout of innervated body structures. The internal layer (red) represents the enteric system. See ***Figure 5— figure supplement 1*** for detailed gustatory receptor expression.

In addition, as we progress from the inner to the outer layers, there is a graded contribution of connections having monosynaptic sensory-to-output contacts (highest being between the inner layers). In other words, the greatest number of monosynaptic connections occur between the enteric system and the neuroendocrine system, followed by the pharyngeal sensory organs to the AN motor neurons, and the least from the external organs. We point out, for example, that the olfactory projections from the external dorsal organ have no monosynaptic connections whatsoever to any output neurons. In this context, the Se0 serotonergic neurons appear to play a special role, as these have the greatest number of monosynaptic contacts from both the enteric system and the pharyngeal sensory organs.

### Multi-synaptic connections to the mushroom body (MB) associative memory circuits

As a contrast to direct input-to-output connections, we additionally looked at connections to a higher brain center for learning and memory, the mushroom body. To this end, we checked previ-ously described projection neurons to the MB calyx (*Eichler et al. 2017*) for inputs from the sensory neurons identified here. Remarkably, the monosynaptic reflex circuit and the multi-synaptic MB projections utilize almost completely different set of sensory synaptic compartments (***Figure 6A***). The three compartments that comprise the vast majority of the monosynaptic circuits (ACa, AVa and AVp) have no outputs onto the MB input projection neurons; rather, three new synaptic compartments are utilized (ACp, ACal and ACpl; ***Figure 6A,B***). Aside from the AL (from which over 20% of output synapses of the ORNs target the MB calyx via olfactory projection neurons), the most prominent synaptic compartment is the ACal, from which almost 40% of the synapses output onto thermosensory projection neurons. We also note that around 45% of all incoming synapses onto the gustatory projections neurons derive from ACp (***Figure 6C***, right panel). This is also consistent with the view that the ACp is the primary sensory compartment onto which the external and pharyngeal gustatory sensory organs project (*Colomb et al. 2007; Hartenstein et al. 2017*).

**Figure 6.**
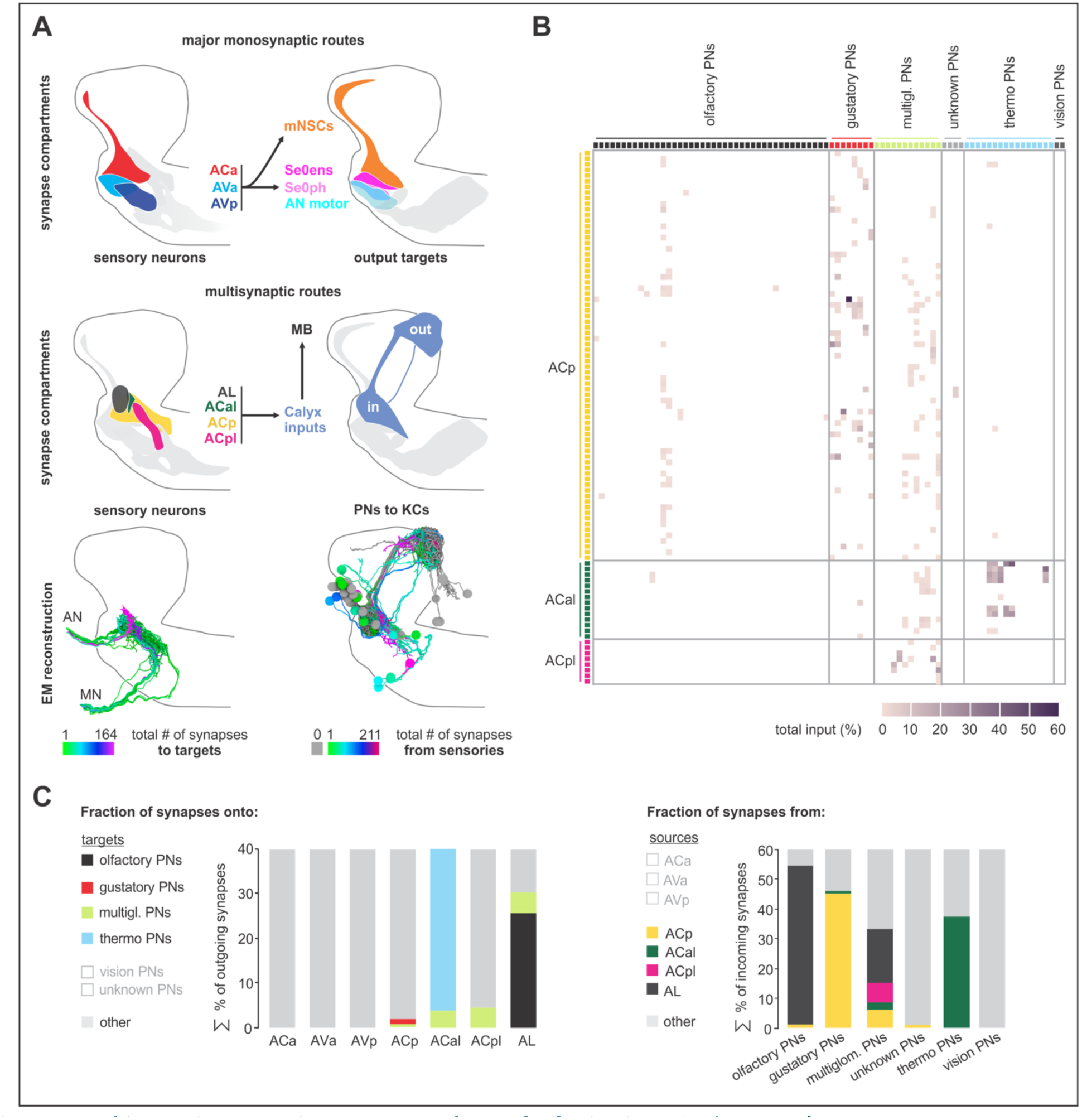
Multisynaptic sensory inputs onto mushroom body circuits. **(A)** Schematic of major monosynaptic routes (top panel). Connectivity between presynaptic sensory neurons (ACa, AVa and AVp) and postsynaptic outputs (mNSCs, Se0ens, Se0ph and AN motor). Schematic of multisynaptic routes to the mushroom body (middle panel). Connectivity between presynaptic sensory neurons (antennal lobe, ACal, ACp, ACpl) and postsynaptic projection neurons to the calyx. EM reconstruction of respective neurons (lower panel); Left: sensory neurons (olfactory receptor neurons are excluded) are color-coded based on total number of synapses to the projection neurons. Right: projection neurons are color-coded based on total number of synapses from sensory neurons. Lateral views show neurons of the right brain hemisphere. **(B)** Adjacency matrix showing sensory-to-PN connectivity, color-coded by percentage of inputs on PN dendrites. Or35a-PN is essentially the only olfactory projection neuron that receives multisensory input from non-olfactory receptor neurons of the ACp (primary gustatory center). **(C)** Left: percentage of presynapses of sensory neurons to PNs. Right: percentage of postsynapses of PNs from sensory neurons.

### Integration of polysynaptic connections onto monosynaptic circuits

We then asked how the hugin neuropeptide (*Drosophila* neuromedin U homolog) circuit, which relays gustatory information to the protocerebrum (*Schlegel et al. 2016; Hückesfeld et al. 2016*), would be positioned with respect to the monosynaptic reflex and multisynaptic MB memory circuits. Based on our earlier studies on mapping sensory inputs onto hugin protocerebrum neurons (huginPC) (*Schlegel et al. 2016; Hückesfeld et al. 2016*), we were expecting most inputs from the ACp, which is the primary gustatory sensory compartment (*Colomb et al. 2007; Hartenstein et al. 2017*). However, most of the huginPC neurons receive inputs from the sensory compartments ACa and AVa, which are the two major monosynaptic compartments that originate from enteric organs. HuginPC neurons do receive inputs from the external and pharyngeal organs (i.e., through sensory compartment ACp), but to a much smaller degree (***Figure 7–figure supplement 1***). Thus, unlike the MB circuit that utilizes a completely new set of sensory inputs, the huginPC circuit is associated with a feeding related monosynaptic circuit.

Based on these observations from the hugin neuropeptide circuit in interconnecting sensory and neuroendocrine outputs, we asked a broader question concerning input-output connections: for any given pair of neurons comprising the monosynaptic reflex circuit, how many additional polysynaptic paths exist and what could be the functional significance of such parallel pathways (*Leonardo 2005*)? To illustrate, we selected a target neuron from each of the three clusters of neurosecretory output cells (Dilps, DMS and DH44) and listed all sensory neurons that make monosynaptic connections with at least 2 synapses (***Figure 7A,B***). We then asked, using the same threshold, how many different di-synaptic paths exist and how often a particular interneuron is used for the different possible converging paths (“degree” of convergence). We also calculated the relative synaptic strengths of the connection among the various paths (“ranking index” of 1.0 represents highest synaptic strength from all possible inputs to the output neuron). Several properties are revealed: (1) different sensory neurons make monosynaptic contacts to a common output target (2) each output neuron can be reached from a given sensory neuron by multiple routes through the use of different interneurons (3) a given interneuron can receive inputs from different sensory neurons to target the same output neuron; this would fit the definition of the “common path” that Sherrington described (*Sherrington 1906*). These observations hold true for the majority of monosynaptic sensory-output pairs we have examined.

A potential functional consequence of such circuit architecture can be seen if we now include all the sensory inputs onto the interneurons. As an example, we take Dilp 1L as the common output, and the interneurons H1 (a huginPC neuron) and “S” (not previously described) as two of the polysynaptic paths onto a common output (***Figure 7C***). One consequence of such superimposition is that the type of sensory information that can reach a common output neuron can be significantly increased. In this case, the Dilp 1L neuron, in addition to receiving inputs from 4 sensory neurons (from monosynaptic paths), now receives inputs from 7 new sensory neurons through the interneuron “S”, and 8 new sensory neurons from interneuron H1. Furthermore, these additional sensory neurons derive from new peripheral regions (e.g., pharyngeal in addition to enteric). Note also that the two interneurons themselves interact, thus increasing the number of paths available between any sensory-output pair. The interneurons could also sharpen sensory information by inhibiting parallel pathways, e.g. through feed-forward inhibition. Extending this to the full set of neurosecretory output targets illustrates the increase in the sensory neuron number and in the peripheral origin that can be gained by integrating polysynaptic connections onto monosynaptic reflex circuits (***Figure 7D***; ***Figure 7–figure supplement 2***): monosynaptic paths would only allow sensory inputs from enteric origin, whereas polysynaptic paths would allow sensory inputs from enteric, pharyngeal and external origins.

**Figure 7.**
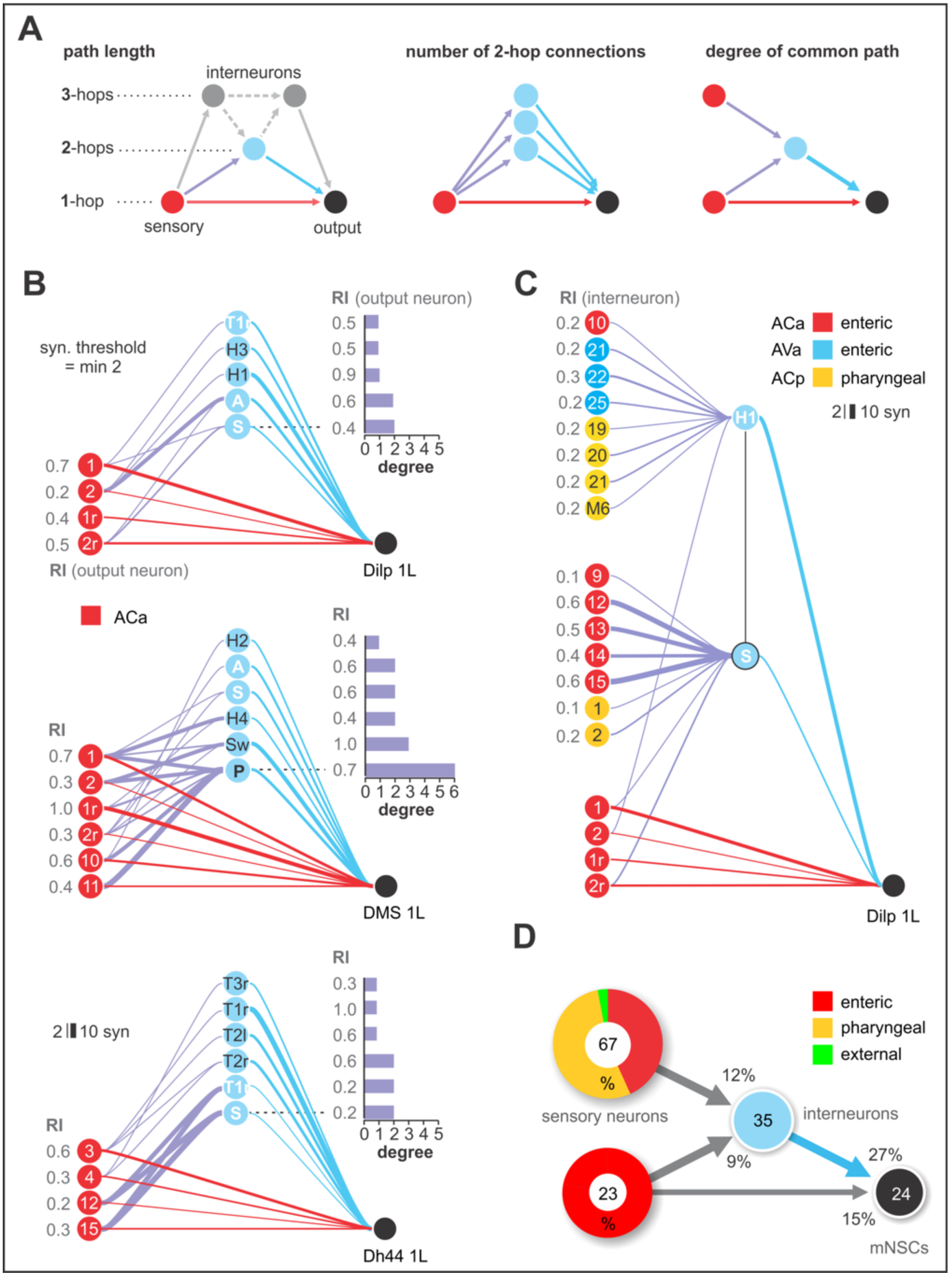
Integration of polysynaptic connections onto monosynaptic circuits. **(A)** Illustration of direct (1-hop) sensory to output neuron connections and indirect (2-hop, 3-hop) paths which involve 1 or 2 interneurons to reach the same output neuron (left panel). Illustration of sensory divergence, which defines the number of possible paths to reach the same target neuron through different interneurons (middle panel). Illustration of sensory convergence, which defines how often (degree) a common path is used by different sensory neurons to reach the same output neuron (right panel). **(B)** All 1- and 2-hop connections for one cell from each of the three clusters of neurosecretory output cells (Dilps, DMS and DH44), using a synaptic threshold of 2. Ranking index (**RI**) shows the relative synaptic strength of every connection compared to the total synaptic input of the output neuron (1.0 represents the highest from all possible inputs to the output neuron). **(C)** Representation of 2 selected alternative paths (“S” and “Hugin-PC left 1”) and their sensory inputs, using a synaptic threshold of 2. Ranking index (**RI**) shows the relative synaptic strength of every connection compared to the total synaptic input of the 2 interneurons. Both interneurons integrate a completely different set of sensory neurons from different sensory compartments to connect onto the basic sensory-to-Dilp circuit. In addition, the Hugin neuron acts as part of a 3-hop connection. **(D)** Summarizing representation of all monosynaptic sensory-to-mNSCs connections (grouped targets: Dilps, DMS, DH44), and their alternative paths through interneurons to reach one cell of the target group, using a synaptic threshold of 2. Note that all alternative paths (interneurons) receive monosynaptic input from 67 other sensory neurons (synaptic threshold= 2), thus integrating a completely different set of sensory neurons onto the basic reflex circuit. Numbers within circles represent number of neurons. Percentages represent fraction of synapses from upstream neurons (arrows). Percentage sensory composition (the two left donut circles) is shown by peripheral origin (enteric, pharyngeal, external). See ***Figure 6–figure supplement 1-2*** for detailed path number and connectivity.

In summary, we propose that the different path possibilities allow different strength and combination of sensory inputs to be evaluated, which would then determine which synaptic path will dominate to a given output. Such multisensory integration via multiple parallel pathways would be necessary to make sense of a complex, multimodal world, and to better choose a behavioral response.

## Discussion

We provide a comprehensive synaptic map of the sensory and motor output neurons that underlie food intake and metabolic homeostasis in *Drosophila* larva. Seven topographically distinct sensory compartments, based on modality and peripheral origin, subdivide the SEZ, a region with functional similarities to the vertebrate brainstem. Sensory neurons that form monosynaptic connections are mostly of enteric origin, and are distinct from those that form multisynaptic connections to the mushroom body (MB) memory circuit. Different polysynaptic connections are superimposed on the monosynaptic sensory-motor pairs that comprise the reflex arc. Such circuit architecture may be used for controlling feeding reflexes and other instinctive behaviors.

### Elemental circuit for feeding

Reflex circuits represent a basic circuit architecture of the nervous system, whose anatomical and physiological foundations were laid down by Cajal and Sherrington (*Ramon y Cajal 1894; Sherrington 1906; Swanson 2011*).The *Drosophila* larval feeding reflex circuit comprises the motor neurons that innervate the muscles involved in pharyngeal pumping, as well as the neurosecretory neurons that target the endocrine organs. They also include a cluster of serotonergic neurons that innervate the entire enteric nervous system, and which may have neuromodulatory effects on the feeding system in a global manner. The vast majority of output neurons are targeted monosynaptically from a set of topographically distinct sensory synaptic compartments in the CNS. These compartments target the output neurons in overlapping domains: the first, ACa, targets all neuroendocrine cells as well as the serotonergic neurons; the second, AVa, targets a subset of neuroendocrine cells, the serotonergic neurons and most of the pharyngeal motor neurons, while the third, AVp, targets the serotonergic neurons and a different set of pharyngeal motor neurons. With these motor outputs, one can in principle fulfill the most basic physiological and behavioral needs for feeding: neurosecretory cells for metabolic regulation and pharyngeal motor neurons for food intake. This set of monosynaptic connections can thus be seen to represent an elemental circuit for feeding, since the connections between the input and output neurons cannot be broken down any further. Vast majority of the sensory inputs comprising this “elemental feeding circuit” derive from the enteric nervous system to target the pharyngeal muscles involved in food intake and neuroendocrine output organs. However, there is a small number of monosynaptic reflex connection that originate from the somatosensory compartment. The output neurons targeted by these somatosensory neurons are motor neurons that control mouth hook movements and head tilting, movements which are involved in both feeding and locomotion. In this context, it is noteworthy that monosynaptic reflex connections are found to a much lesser degree in the larval ventral nerve cord, which generates locomotion (unpublished data from *Ohyama et al., 2015*). An analogous situation exists in *C. elegans*, where majority of the monosynaptic reflex circuits are found in the head motor neurons and not in the body (*Yan et al. 2017*). One reason could be due to the relative complexity in the response necessary for food intake as compared to locomotion. For example, a decision to finally not to swallow a harmful substance, once in the mouth, may require a more local response, e.g. muscles limited to a very specific region of the pharynx and esophagus, where monosynaptic arc might suffice. By contrast, initiating escape behaviors requires a more global response with respect to the range and coordination of body movements involved, although it also employs multimodal sensory integration via a multilayered circuit (*Ohyama et al. 2015*).

### Monosynaptic connections between the sensory neurons

The inter-sensory connections show a combination of hierarchical and reciprocal connections, which may increase the regulatory capability and could be especially important for monosynaptic circuits. By contrast, very few monosynaptic connections exist between the larval olfactory, chordotonal or nociceptive class IV sensory in the body (*Ohyama et al. 2015; Jovanic et al. 2016; Gerhard et al. 2017*). Interestingly, there is also a much higher percentage of intersensory connections between olfactory neurons in the adult as compared to the larva, which could function in gain modulation at low signal intensities (*Tobin et al. 2017*). This might be attributable to adults requiring faster processing of olfactory information during flight navigation (or mating), and/or to minimize metabolic cost (*Wilson 2014*). Whether such explanation also applies to the differences in intersensory connection between the different types of sensory neurons in the larvae remains to be determined.

### Superimposition of polysynaptic pathways onto monosynaptic circuits

We found very few cases where a monosynaptic path between any sensory-output pair is not additionally connected via a polysynaptic path. An interesting question in the context of action selection mechanism is which path a sensory signal uses to reach a specific target neuron. For example, a very strong sensory signal may result in a monosynaptic reflex path being used. However, a weaker sensory signal may result in using a different path, such as one with less threshold for activation. This would also enable the integration of different types of sensory signals through the usage of multiple interneurons, since the interneurons may receive sensory inputs that are not present in monosynaptic connections. For example, sensory neurons can target the neuroendocrine cells directly (monosynaptically), or through a hugin interneuron (disynaptically). The sensory compartment that directly target the neuroendocrine cells are of enteric origin; however, when hugin neurons are utilized as interneurons, not only is the number of sensory neurons from the same sensory compartment increased, but sensory neurons are added from a completely new peripheral origin. Thus, the hugin interneurons enable sensory inputs from different peripheral origins, e.g. to integrate enteric inputs with pharyngeal gustatory inputs, to influence an output response, which, in this case, is to stop feeding (*Schoofs et al. 2014a*).

The coexistence of polysynaptic and monosynaptic paths could also be relevant for circuit variability and compensation (*Leonardo 2005; Marder and Goaillard 2006*): destruction of any given path would still enable the circuit to function, but with more restrictions on the precise types of sensory information it can respond to. In certain cases, this may even lead to strengthening of alternate paths as a form of synaptic plasticity.

An open issue is how the sensory synaptic compartments might be connected to the feeding central pattern generators (CPGs) which have been demonstrated to exist in the SEZ (*Schoofs et al. 2010; Hückesfeld et al. 2015*), especially since CPGs are defined as neural circuits that can generate rhythmic motor patterns in the absence of sensory input. However, the modulation of CPG rhythmic activity can be brought about by sensory and neuromodulatory inputs (*Marder and Bucher 2001; Marder 2012*). A complete circuit reconstruction of the larval SEZ circuit may shed some light on the circuit structure of feeding CPGs.

### Multisynaptic sensory inputs onto memory circuits

A more complex circuit architecture is represented by the MB, the site of associative learning and memory in insects: a completely different set of sensory synaptic compartments is used to connect the various projections neurons to the MB calyx. Thus, the MB module is not superimposed onto the monosynaptic reflex circuits but rather forms a separate unit. The outputs of the MB calyx remains largely unknown, but they must at some point connect to the motor or neuroendocrine neurons. The classical studies by Pavlov demonstrated conditioned reflex based on an external signal and an autonomic secretory response in response to food (*Pavlov 1927; Todes 2001*). Although a comparable autonomic response has not been analyzed in the larvae, analogous associative behavior based on odor choice response has been well studied (*Aceves-Pina and Quinn 1979; Gerber and Stocker 2007; Eichler et al. 2017; Widmann et al. 2018*). It is also noteworthy that in the *Aplysia*, classical conditioning of the gill withdrawal reflex involves monosynaptic connections between a sensory neuron (mechanosensory) and a motor neuron, and neuromodulation by serotonin (*Bailey et al. 2000*). This constellation has similarities with the elemental feeding circuit consisting of sensory, motor and serotonergic modulatory neurons. For more complex circuits of feeding behavior in the mouse, a memory device for physiological state, such as hunger, has been reported involving synaptic and neuropeptide hormone circuits (*Yang et al. 2011*). Identification of output neurons of the MB calyx would help address how memory circuits interact with reflex feeding circuits.

### Control of reflexes

Feeding behavior manifests itself from the most primitive instincts of lower animals, to deep psychological and social aspects in humans. It encompasses cogitating on the finest aspects of food taste and the memories evoked by the experience, to sudden reflex reactions upon unexpectedly biting down on a hard seed or shell. Both of these extremes are mediated, to a large degree, by a common set of feeding organs, but the way these organs become utilized can vary greatly. The architecture of the feeding circuit described here allows the various types of sensory inputs to converge on a limited number of output responses. The monosynaptic pathways would be used when fastest response is needed. The presence of polysynaptic paths would enable slower and finer control of these reflex events by allowing different sensory inputs, strengths or modalities to act on the monosynaptic circuit. This can be placed in the context in the control of emotions and survival circuits (*LeDoux 2012*), or by cortex regulation of basic physiological or autonomic processes (*Dum et al. 2016*). In a striking example, pupil dilation, a reflex response, has been used as an indicator of cognitive activity (*Hess and Polt 1964; Kahneman and Beatty 1966; Larsen and Waters 2018*). Here, a major function of having more complex circuit modules on top of monosynaptic circuits may be to allow a finer regulation of feeding reflexes, and perhaps of other reflexes or instinctive behaviors.

As an outlook, our analysis provides an architectural framework of how a feeding circuit is organized in the CNS. The circuit is divided into two main axes that connect the input to the output systems: the sensory-neurosecretory cell axis and the sensory-motor neuron axis (*Swanson 2011*). The sensory system targets overlapping domains of the output neurons; for example, a set of sensory neurons targets exclusively the neuroendocrine cells, other targets both neuroendocrine and pharyngeal motor neurons, and another just the pharyngeal motor neurons. The inputs derive mostly from the internal organs. These connections form the monosynaptic reflex circuits. With these circuits, one can perform the major requirements of feeding regulation, from food intake and ingestion to metabolic homeostasis. Additional multi-synaptic circuits, such as the CPGs, those involving sensory signaling from the somatosensory system (external inputs), or those comprising the memory circuits, are integrated or added to expand the behavioral repertoire of the animal (***Figure 8***). Although circuit construction may proceed from internal to the external, the sequence is reversed in a feeding animal: the first sensory cues are external (olfactory), resulting in locomotion (somatic muscles) that can be influenced by memory of previous experience; this is followed by external taste cues, resulting in food intake into the mouth; the final action is the swallowing of food, involving pharyngeal and enteric signals and reflex circuits. However, regardless of the types of sensory inputs, and whether these are transmitted through a reflex arc, a memory circuit or some other multi-synaptic circuits in the brain, they will likely converge onto a certain set of output neurons, what Sherrington referred to as the “final common path” (*Sherrington 1906*). The current work is a first step towards finding the common paths.

**Figure 8.**
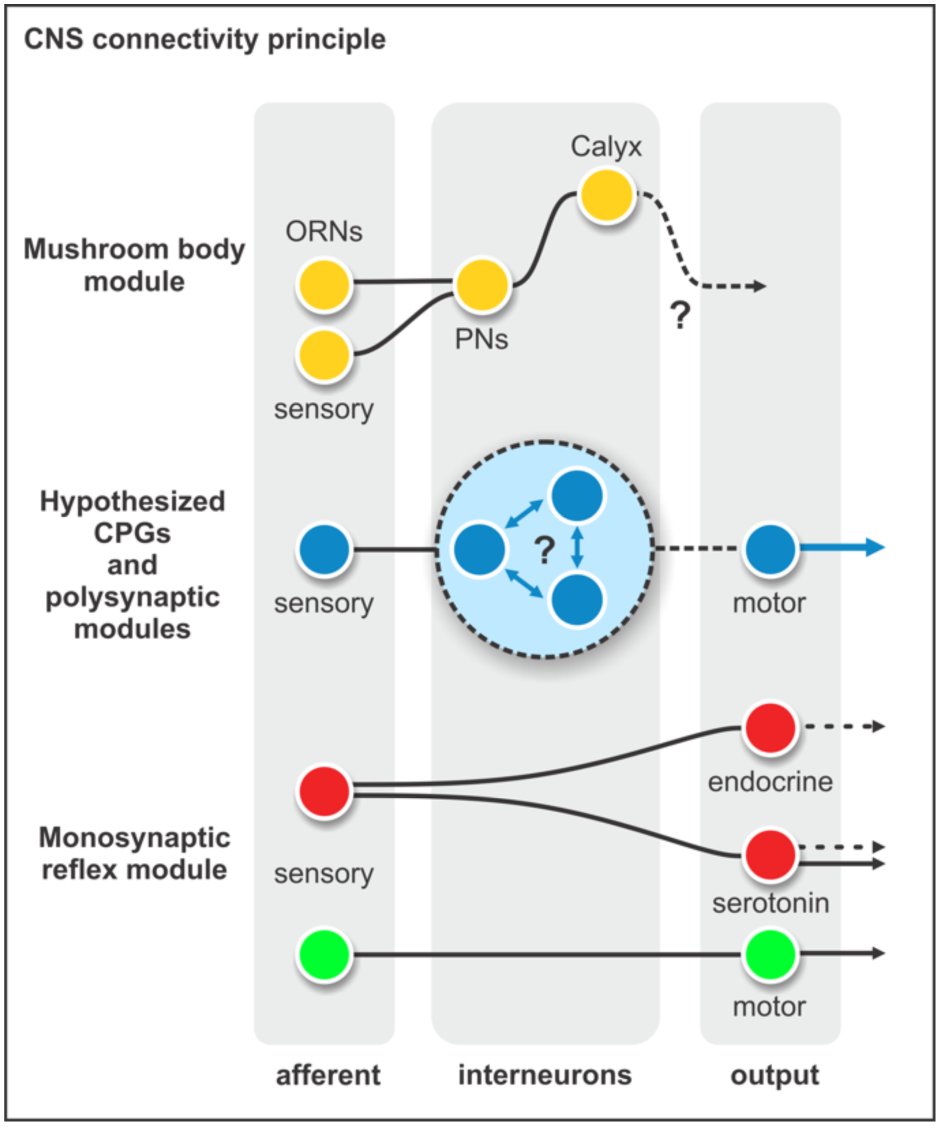
Connectivity principles in the brain. Different poly synaptic modules are integrated onto existing monosynaptic circuits, or added separately as new multi-synaptic circuits, e.g. the mushroom body module and the hypothesized CPG-module.

## Materials and methods

### Neuronal reconstruction

All reconstructions were based on an ssTEM (serial section transmission electron microscope) data set of a complete nervous system of a 6-h-old [iso] CantonS G1 x w1118 larva as described in (*Ohyama et al. 2015*). Using a modified version of the web-based software CATMAID (*Saalfeld et al. 2009*) we manually reconstructed neurons’ skeletons and annotated synapses following the methods described in (*Ohyama et al. 2015*) and (*Schneider-Mizell et al. 2016*). Sensory and motor neurons were identified by reconstructing all axons in the antennal nerve, maxillary nerve and the prothoracic accessory nerve. Further, neurons with their soma in the brain and projections through one of the three pharyngeal nerves have been identified as motor neurons and serotonergic output neurons. All annotated synapses represent fast, chemical synapses equivalent to previously described typical criteria: thick black active zones, preand postsynaptic membrane specializations (*Prokop and Meinertzhagen 2006*).

### Morphology similarity score

To neuron morphologies (***Figure 1; Figure 1–figure supplement 2-4; Figure 2–figure supplement 6***), we used a morphology similarity score described by *Kohl et al. 2013*. Briefly, reconstructions of neurons are converted to “dotprops”, 3d positions with an associated tangent vector: for each dotprop from a query neuron, the closest point on a target neuron was determined and scored by distance and the absolute dot product of their two tangent vectors. The total similarity score is the average score over all point pairs between query neuron Q and target neuron T:

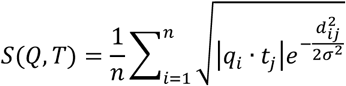

where n is the number of points in the query neuron, d_ij is the distance between point i in the query neuron and its nearest neighbor, point j, in the target neuron and q_i and t_j are the tangent vectors at these points. σ determines how close in space points must be to be considered similar. For our calculations, we used σ of 2 um. Similarity score algorithm was implemented in a Blender plugin (https://github.com/schlegelp/CATMAID-to-Blender).

### Synapse similarity score

To calculate similarity of synapse placement between two neurons, we calculated the synapse similarity score (***Figure 2; Figure 2–figure supplement 1-6; Figure 3–figure supplement 3-5***):

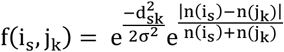

With the overall synapse overall synapse similarity score for neurons i and j being the average of f(i_s_, j_k_) over all synapses s of i. Synapse k being the closest synapse of neuron j to synapse s [same sign (pre-/post-synapse) only]. d_sk_ being the linear distance between synapses s and k. Variable σ determines which distance between s and k is considered as close. n(j_k_) and n(i_s_) are defined as the number of synapses of neuron j/i that are within a radius ω of synapse k and s, respectively (same sign only). This ensures that in case of a strong disparity between n(i_s_) and n(j_k_), f(i_s_, j_k_) will be close to zero even if distance d_?A_ is very small. Values used: σ = ω = 2000 nm. Similarity score algorithm was implemented in a Blender plugin (https://github.com/schlegelp/CATMAID-to-Blender).

### Normalized connectivity similarity score

To compare connectivity between neurons (***Figure 2–figure supplement 6***), we used a modified version of the similarity score described by *Jarrell et al., 2012*:

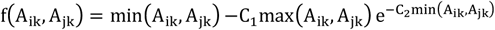

With the overall connectivity similarity score for vertices i and j in adjacency matrix A being the sum of f(A_ik_, A_jk_) over all connected partners k. C_1_ and C_2_ are variables that determine how similar two vertices have to be and how negatively a dissimilarity is punished. Values used were: C_1_ = 0.5 and C_2_ = 1. To simplify graphical representation, we normalized the overall similarity score to the minimal (sum of −C_1_max(A_ik_, A_jk_) over all k) and maximal (sum of max(A_ik_, A_jk_) over all k) achievable values, so that the similarity score remained between 0 and 1. Self-connections (A_ii_, A_jj_) and A_ij_ connections were ignored.

### Clustering

Clusters for dendrograms were created based on the mean distance between elements of each cluster using the average linkage clustering method. Clusters were formed at scores of 0.1 for synapse similarity score (***Figure 2; Figure 2–figure supplement 1-5***).

### Percentage of synaptic connections

Percentage of synaptic connections was calculated by counting the number of synapses that constitute between neuron A and a given set of pre- or postsynaptic partners divided by the total number of either incoming our outgoing synaptic connections of neuron A. For presynaptic sites, each postsynaptic neurite counted as a single synaptic connection (***Figure 3; Figure 3–figure supplement 3-5; Figure 4; Figure 6; Figure 7–figure supplement 1***).

### Ranking index

Ranking index was calculated by counting the number of synapses that constitute between neuron A and a given target neuron B divided by the highest number of synapses among all incoming synaptic connections of target neuron B(***Figure 7; Figure 7–figure supplement 2***).

### Neuronal representation

Neurons were rendered with Blender 3D (ww.blender.org) and edited in Adobe Corel Draw X7 (www.corel.com). A script for a CATMAID-Blender interface is on Github (https://github.com/schlegelp/CATMAID-to-Blender)

### Graphs

Graphs were generated using Excel for Mac v15.39 (www.microsoft.com), Sigma Plot 12.0 (www.sigmaplot.com) and edited in Corel Draw X7.

### Flies

The following GAL4 driver and UAS effector lines were used: Gr2a-GAL4 (Bloomington #57589), Gr10a-GAL4 (Bloomington #57597), Gr22b-GAL4 (Bloomington #57604), Gr22e-GAL4 (Bloomington #57608), Gr23a-GAL4 (Bloomington #57611), Gr28a-GAL4 (Bloomington #57614 and #57613), Gr28b-GAL4 (*Scott et al. 2001*), Gr28b.a-GAL4 (Bloomington #57615), Gr28b.e-GAL4 (Bloomington #57621), Gr32a-GAL4 (Bloomington #57622), Gr33a-GAL4 (Bloomington #57624), Gr39a.a-GAL4 (Bloomington #57631), Gr39a.b-GAL4 (Bloomington #57632), Gr39a.d-GAL4 (Bloomington #57634), Gr39b-GAL4 (Bloomington #57635), Gr43a-GAL4 (Bloomington #57636 and #57637), Gr43aGAL4 knock-in (*Miyamoto et al. 2012*), Gr57a-GAL4 (Bloomington #57642), Gr58b-GAL4 (Bloomington #57646), Gr59a-GAL4 (Bloomington #57648), Gr59d-GAL4 (Bloomington #57652), Gr63a-GAL4 (Bloomington #57660), Gr66a-GAL4 (*Scott et al. 2001*), Gr68a-GAL4 (Bloomington #57671), Gr77a-GAL4 (Bloomington #57672), Gr93a-GAL4 (Bloomington #57679), Gr93b-GAL4 (Bloomington #57680), Gr93c-GAL4 (Bloomington #57681), Gr93d-GAL4 (Bloomington #57684), 94a-GAL4 (Bloomington #57686), Orco-GAL4 (Bloomington #23909) and 10X-UAS-mCD8::GFP (Bloomington #32184)

### Immunohistochemistry

Dissected larval brains with attached CPS and intact pharyngeal nerves were fixed for 1 hr in paraformaldehyde (4%) in PBS, rinsed with PBS-T and blocked in PBS-T containing 5% normal goat serum. For antibody stainings of Gr-GAL4>10xUAS-mCD8::GFP primary antibody were conjugated goat anti-GFP (1:500, Abcam, ab6662), mouse anti-fasciclin2 (1:500, DSHB) and mouse anti-22C10 (1:500, DSHB) and the secondary antibody was anti-mouse Alexa Flour 568 (1:500, Invitrogen). Brains were rinsed with PBS-T and mounted in Mowiol (Roth, 0713). For antibody stainings of Gr43a-GAL4>10xUAS-mCD8::GFP primary antibody were rabbit anti-GFP (1:500, Abcam, ab6556), mouse anti-fasciclin2 (1:500, DSHB) and mouse anti-22C10 (1:500, DSHB) and the secondary antibody were anti-rabbit Alexa Flour 488 (1:500, Invitrogen) and anti-mouse Alexa Flour 568 (1:500, Invitrogen). Brains were rinsed with PBS-T and dehydrated through an ethanol-xylene series and mounted in DPX. Imaging was carried out using a Zeiss LSM 780 confocal microscope with a 25x objective (Zeiss).

## Acknowledgements

We thank Sarah Brenner, Benjamin White, Henning Fenselau and William Schafer for their timely contributions, and the Bloomington Stock Center for fly strains. We also thank all the “tracers” for their contribution to the EM reconstruction.

## Additional information

### Funding

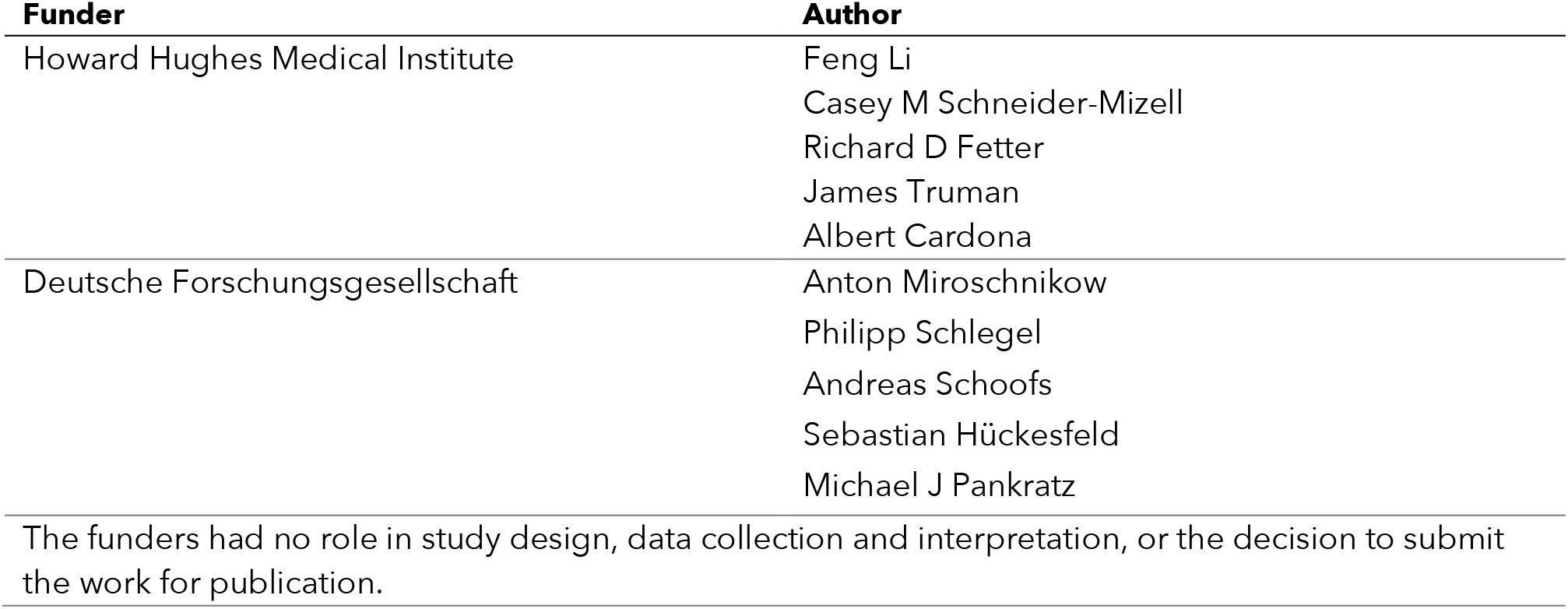

## Author contributions

AM, PS, AC, Conception and design, Acquisition of data, Analysis and interpretation of data, Drafting or revising the article; AS, SH, FL, Acquisition of data; CMS-M, RDF, JWT, Generation of dataset, Drafting or revising the article; MJP, Conception and design, Analysis and interpretation of data, Drafting or revising the article.

## Additional files

### Supplementary files

Supplementary file 1.

